# Spatio-temporal dynamics of face perception

**DOI:** 10.1101/550038

**Authors:** I. Muukkonen, K. Ölander, J. Numminen, V.R. Salmela

## Abstract

The temporal and spatial neural processing of faces has been investigated rigorously, but few studies have unified these dimensions to reveal the spatio-temporal dynamics postulated by the models of face processing. We used support vector machine decoding and representational similarity analysis to combine information from different locations (fMRI), time windows (EEG), and theoretical models. By correlating information matrices derived from pairwise classifications of neural responses to different facial expressions (neutral, happy, fearful, angry), we found early EEG time windows (starting around 130 ms) to match fMRI data from early visual cortex (EVC), and later time windows (starting around 190ms) to match data from occipital and fusiform face areas (OFA/FFA) and posterior superior temporal sulcus (pSTS). According to model comparisons, the EEG classifications were based more on low-level visual features than expression intensities or categories. In fMRI, the model comparisons revealed change along the processing hierarchy, from low-level visual feature coding in EVC to coding of intensity of expressions in the right pSTS. The results highlight the importance of a multimodal approach for understanding the functional roles of different brain regions in face processing.

## 1 Introduction

Faces contain information about different socially important categories, such as identity and the emotional state of an individual. Behaviorally, we excel at distinguishing these different sources of information, and can easily identify a familiar person just from their face despite changes in expression or viewpoint. Indeed, the processing of faces has been suggested to be ‘special’ (Richler & Gauthier, 2014), and faces have been proposed to be processed more holistically than other visual stimuli (Shen & Palmeri, 2015; Tanaka & Simonyi, 2016; but see also Gold, Mundy, & Tjan, 2012). Newborns recognize their mother’s face (Bushnell, 2001; Bushnell, Sai, & Mullin, 1989) despite their otherwise undeveloped visual skills (Dobson & Teller, 1978), fetuses focus more on face-shaped than non-face-shaped objects (Reid et al., 2017) and inversion distorts face recognition more than recognition of other objects (Taubert, Apthorp, Aagten-Murphy, & Alais, 2011; Valentine, 1988; Yin, 1969). Thus, faces provide an excellent case for studying how our brain processes nuanced multidimensional information, in which the change in one dimension (e.g. expression) must be separated from the changes in another dimension (e.g. identity).

Several models explaining the neural processing of faces have been proposed (Duchaine & Yovel, 2015; Haxby, Hoffman, & Gobbini, 2000). One main feature in these models is the separation of the processing of changeable and invariant aspects. Changeable aspects, (or motion: Bernstein & Yovel, 2015) such as expressions, are mainly processed in the dorsal stream, especially in the superior temporal sulcus (STS; Greening, Mitchell, & Smith, 2018; Said, Haxby, & Todorov, 2011; Zhang et al., 2016). In contrast, invariant aspects, such as identity, are processed in the ventral stream from part-based processing in the occipital face area (OFA; Atkinson & Adolphs, 2011; Henriksson, Mur, & Kriegeskorte, 2015; Pitcher, Walsh, & Duchaine, 2011), to the fusiform face area (FFA; Anzellotti, Fairhall, & Caramazza, 2014; Carlin & Kriegeskorte, 2017; Dobs, Schultz, Bülthoff, & Gardner, 2018; Kanwisher & Yovel, 2006), and finally to highest-level, viewpoint-invariant processing in the ventral anterior temporal lobe (vATL; Anzellotti & Caramazza, 2016; Anzellotti et al., 2014; Collins & Olson, 2015; Kriegeskorte, Formisano, Sorger, & Goebel, 2007). Hierarchical processing is supported by single-cell recordings from macaques that have found viewpoint-specific coding of face identities in the middle lateral and middle fungus, mirror-symmetrical processing in the anterior lateral patch, and finally almost viewpoint-invariant identity representations in the anterior medial patch (Chang & Tsao, 2017; Freiwald & Tsao, 2010).

Said and colleagues (2010) found the neural activity patterns elicited by video clips of expressions to be similar to behavioral similarity ratings of the expressions in STS, suggesting the processing of changeable features there. Processing in the STS has also been studied by contrasting different (full) expressions to each other or to neutral faces. Specifically, Greening and colleagues (2018) showed significant cross-classification of happy and neutral faces from other expressions when training with eyes and testing on faces with eyes excluded, and vice versa. However, they found no significant cross-classification of fearful, angry, and disgusted faces. Similarly, Zhang and colleagues (2016) found STS to best separate neutral faces from fearful, angry and happy faces. The STS thus seems to be especially sensitive to differences between expressional and neutral faces.

Temporally, the processing of faces starts around 90 ms (Dima, Perry, Messaritaki, Zhang, & Singh, 2018; Sugase, Yamane, Ueno, & Kawano, 1999), perhaps already being somewhat sensitive to differences in expression (Dima et al., 2018; Müller-Bardorff et al., 2018). Most robust evidence for separating faces from other objects is found in the N170-component (at ~170 ms; for a review, see Rossion & Jacques, 2011), which is also sensitive to some expressions (Hinojosa, Mercado, & Carretié, 2015). Coding of identities is usually found later. Although earliest results are found from ~100 ms onwards (Ambrus, Kaiser, Cichy, & Kovács, 2019; Vida, Nestor, Plaut, & Behrmann, 2017), higher-level identity representations surpassing low-level visual information (Vida et al., 2017), as well as sensitivity to familiar faces (Schweinberger & Neumann, 2016), are found later, at around 250 ms, and differentiation of same sex identities only after 400 ms (Ambrus et al., 2019).

Although much is known about the face-specific brain regions and the temporal time course of face processing, a challenge still exists regarding how to combine our understanding of the spatial and temporal aspects of face processing. One possibility is to compare response magnitudes captured with one method to those captured with another method. In a simultaneous EEG-fMRI study, face selectivity (response magnitude between faces and images of chairs) in fMRI was found to correlate with earlier EEG timepoints in OFA than in FFA and STS (Sadeh, Podlipsky, Zhdanov, & Yovel, 2010). Another possibility to combine different imaging methods is to use multivariate analysis methods (Edelman, Grill-Spector, Kushnir, & Malach, 1998; Haxby et al., 2001; Kriegeskorte, Mur, & Bandettini, 2008), such as stimulus reconstruction, classification/decoding and representational similarity analysis (RSA; Kriegeskorte et al., 2008), which can capture more nuanced information structures than univariate methods. In RSA, the data are projected to a geometrical space – representational dissimilarity matrix (RDM) – that highlights the relative differences between responses, not the response amplitudes per se. As these RDMs are indifferent to the type of data that they are derived from, the RDM from a certain timepoint in EEG, for example, can be compared to the RDMs from a certain cluster of voxels in fMRI, thus combining the good temporal accuracy of M/EEG with the good spatial accuracy of fMRI. This approach has been used in studying audiovisual attention networks (Salmela, Salo, Salmi, & Alho, 2018), visual object categorization (Cichy & Pantazis, 2017; Cichy et al., 2014, 2016), and differences between neural responses to tasks and images (Hebart, Bankson, Harel, Baker, & Cichy, 2018). These studies showed that, within the first few hundred milliseconds, processing of visual object categories spreads from early visual areas (V1) to the lateral occipital complex (LOC; Cichy et al., 2016; Kietzmann et al., 2019). In the current study, we apply this method to study the processing of faces.

We used RSA to compare neural representations of faces as characterized with EEG and fMRI to reveal the spatio-temporal dynamics of face processing (Figure 1). We compared representational dissimilarity matrixes (RDMs) – based on pairwise decoding analyses– derived from different time windows in the EEG data and from searchlight voxels in the fMRI data. Our stimuli were faces with different identities (2 males, 2 females, 12 identity-morphs) and different expressions, which varied both in their category (neutral, happy, fearful, and angry) and intensity (100% intensity and morphed 50% intensity). We also compared the data to models representing either expression category, expression intensity, or low-level visual features, allowing us to look for possibly different neural codes in different parts and at different times in the brain. While several studies have looked at neural representations of different expression categories, and a few expression intensities (Surguladze et al., 2003; Winston, O’Doherty, & Dolan, 2003), our study provides a novel way to compare whether expression categories and intensities have different representations in the brain. Especially, several studies have found STS to separate particularly neutral faces from expressions (Greening et al., 2018; Zhang et al., 2016). By including morphed expressions, our study design provides an opportunity to look more closely at whether these results were due to expression intensity coding in the STS.

**Figure 1.**
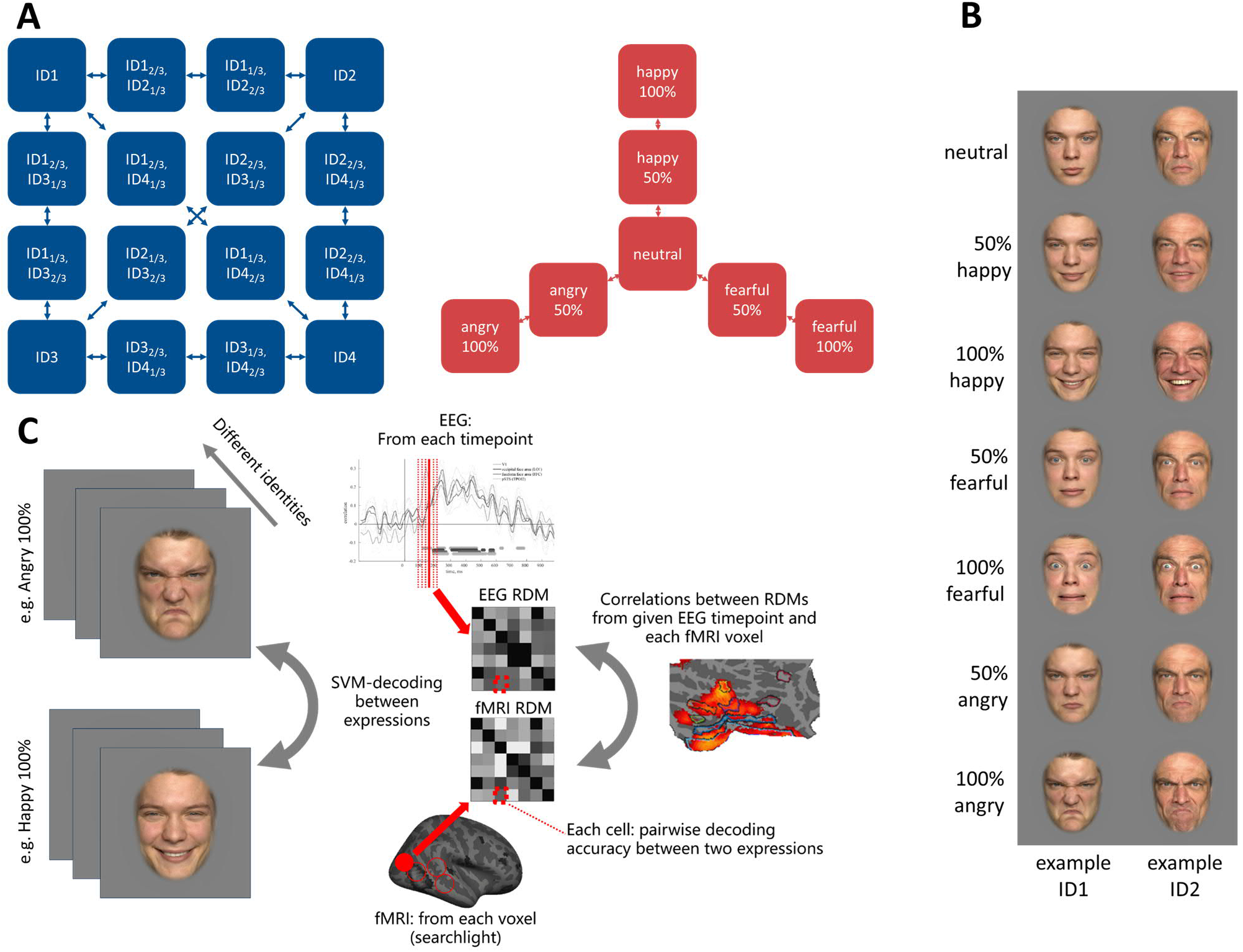
Morph design, example stimuli and analysis design. **A)** Both expressions and identities were morphed. Original images from 4 identities with 4 expressions (neutral, happy, fearful, and angry) were morphed between the identities (left), and between neutral and other expressions within each identity (right). **B)** Stimuli examples of all used expression-morphs from two identities. The example identities are samples from the Faces database (Ebner et al., 2010; https://faces.mpdl.mpg.de/imeji/); different identities were used in the experiment. Identity-morphs are not shown due to copyright restrictions. **C)** Study design for expression EEG-fMRI analyses. Each possible pair of expressions and morphed expressions were classified with SVM, separately in EEG and in fMRI. The different identities were used as exemplars in the classification analyses. From these pairwise classifications, a representational dissimilarity matrix (RDM) was created for each EEG time window and each fMRI searchlight voxel. In the RDMs, each cell is a decoding accuracy between two expression-morphs (e.g. angry 100% vs happy 100%). The RDMs from EEG and fMRI were then correlated with each other, as well as with models.

## 2 Methods

### 2.1 Participants

A total of 18 volunteers were recruited to participate in the study. One participant dropped out before completing the first part. Thus, 17 people (7 males, mean age = 24, SD = 3.4) completed the study. All participants were right-handed, had normal or corrected to normal vision and had no (self-reported) diagnosed difficulties in recognizing faces (e.g. prosopagnosia). They were recruited through the students’ mailing list of the Faculty of Behavioural Sciences at the University of Helsinki, and they received monetary compensation for the participation.

### 2.2 Ethics

The present study was conducted in accordance with the declaration of Helsinki and was reviewed and approved by the Ethics Review Board in the Humanities and Social and Behavioural Sciences of the University of Helsinki. All participants gave informed consent and were screened to be suitable for fMRI scanning with the standard procedure of the AMI centre of Aalto University.

### 2.3 Stimuli

The stimuli consisted of 112 full-colour images of faces. The root-mean-square contrast of the images was normalized to 0.2, and width and height were approximately 8° and 11°, respectively. The images contained four identities (2 females, 2 males) from the Faces database (Ebner, Riediger, & Lindenberger, 2010), and four different expressions for each identity (neutral, angry, happy and fearful). These original images (4 expressions from 4 identities) were first preprocessed with Corel Paintshop Pro X7. The images were 1) straightened horizontally, using the centres of the eyes as a reference line; 2) resized to have the same interocular distance; and 3) centred in a way that the centres of the eyes in each face image were approximately in the same location. This was done to control for irrelevant visual variability in the pictures, for example size or tilt of the face, and to improve face morphing.

After the preprocessing of the original faces, they were morphed with FantaMorph (version 5.4.6). Each original identity with a given expression was morphed to every other identity with the same expression. From these morph dimensions, two morphed images, 1/3 of identity X and 2/3 of identity Y, and vice versa, were used in the study. As a result, there were 12 identity-morphs in addition to the original four identities. Both the original 4 identities and the 12 identity-morphs were then morphed (within an identity) from neutral to the three other expressions, and the 50 % morphs were used in the study (Figure 1A). Thus, the stimulus set contained 112 images in a 16 identity-morphs (4 original, 12 morphed) × 7 expression-morphs (neutral, 3 original-100%, 3 morphed-50%) design. For simplicity, we refer henceforth to both the original and the morphed identities and expressions with the terms identity-morphs and expression-morphs.

### 2.4 Study design

The study contained two parts, fMRI measurement and EEG measurement, which were conducted on different days. Nine of the participants completed the fMRI part first, and eight completed the EEG part first. The time between the measurements was 1 – 40 days for the different participants. Both sessions lasted approximately 1.5 hours, including preparation.

The primary task and stimuli were identical in both measurements. Participants saw a face on a screen (500 ms), and they were instructed to answer, by pressing one of two buttons, whether the face on the screen was female or male. The gender identification task was used in order to ensure participants’ attentional focus on the faces, and it was not trivial, since half of our stimuli were morphs between genders. The inter-stimulus interval was 2500 ms. The study was divided in three runs. Each stimulus was shown twice in each run, once (in random order) before a 30 sec rest period and a second time (in random order) after the rest period. In total, each run contained 224 trials and the duration of each run was approximately 12 minutes. Between the runs the participants were contacted through microphone and were given a rest period if necessary.

The fMRI-measurement began with an anatomical scan (6 min) and ended with a functional face area localizer run. In the localizer scan, intact faces, phase scrambled faces, and checkerboard stimuli were shown while the participant was instructed to fixate at the centre of the screen. In the EEG measurement, video clips of facial expressions were shown at the end of the experiment. The results of these are not, however, discussed here.

### 2.5 Acquisition and preprocessing of EEG

EEG data were recorded using a 64-channel cap and 6 additional electrodes (mastoids, HEOGs from both eyes and two VEOGs from the left eye) with Biosemi Actiview. The EEG data were sampled at 1024 Hz and referenced online to Common Mode Sense (CMS) electrode at PO1.

EEG preprocessing was performed with EEGLAB v13.6.5b running in Matlab R2018a. The data were first downsampled to 200Hz, and bandpass-filtered between 0.1 – 81 Hz (- 6db cutoff points, acausal filter). Line noise was removed with the Cleanline-plugin (Mullen, 2012) and bad channels detected and removed with the clean_rawdata v0.31-plugin (Kothe, 2013). The data were then re-referenced to the average of the 64 cap electrodes, the removed channels were interpolated and all the non-cap electrodes were removed. Epochs of −300 – 1000 ms from stimuli onset were created, and epochs with the same stimuli were averaged. No baseline correction was applied. Thus, a four-dimensional EEG-data-matrix was created, the dimensions being the 17 participants, 112 different stimulus types, 64 channels and 260 timepoints (5 ms each). The 260 timepoints were later collapsed in the decoding analyses into final 130 time windows of 10 ms each (see *2.7. Decoding*).

### 2.6 Acquisition and preprocessing of fMRI

Functional MRI data were recorded using a 3T MAGNETOM Skyra scanner (Siemens Healthcare, Erlangen, Germany) at the AMI-Centre of Aalto University School of Science. A 30-channel head coil was used. The functional echo-planar images (EPI) were acquired with an imaging area consisting of 43 contiguous slices (TR 2.4 s, TE 32 ms, voxel matrix 64 · 64, field of view 20 cm, slice thickness 3.0 mm). Three functional runs of 275 volumes (including 4 initial dummy volumes) were measured, each lasting approximately 12 minutes. In preprocessing, slice timing and motion correction (but no smoothing) were applied, and all images were co-registered to a T1 anatomical image (MPRAGE). Finally, the images from each subject were normalized to the standardized MNI head space. The fMRI data were modelled with an event-related GLM analysis containing separate regressors for each stimulus type (112 in total), and 6 nuisance regressors for motion. As a result, there were 112 (stimuli) × 3 (runs) whole-brain beta images for each subject. fMRI analyses were conducted with the SPM12 toolbox for Matlab.

### 2.7 Decoding analyses

We used leave-one out support vector machine (SVM; Cortes & Vapnik, 1995) decoding with the non-decoded stimulus dimension (expression or identity) as exemplars, and voxels/EEG-channels as features. Thus, two sets of analyses were conducted, one to study identities and one to study expressions. For expressions, a pairwise decoding analysis was conducted between each possible expression-morph pair (e.g. 50% morphed happy vs. neutral), resulting in 21 pairwise decoding analyses for expressions in total (number of possible pairs from 7 different stimuli: 6+5+4+3+2+1). Likewise, for identities, a pairwise decoding analysis was conducted between each possible identity-morph pair (e.g., identityA vs. identity-morphA_1/3_B_2/3_), resulting in 120 pairwise decoding analyses for identities (number of possible pairs from 16 different stimuli: 15+14…+1). In each pairwise decoding analysis, the data were divided into training and test sets, leaving either one of the non-decoded stimulus categories (e.g. one expression-morph when decoding identities) out (EEG), or one run out (fMRI), and this procedure was repeated until every category/run was used as the test set. In EEG, expression decoding analyses were repeated 16 times (leaving one identity-morph out at a time) and identity decoding analyses 7 times (leaving one expression-morph out at a time). Thus, expression decoding analyses contained 15+15 exemplars in the training set and 1+1 in the test set, and identity decoding analyses contained 6+6 exemplars in the training set and 1+1 in the test set.

In fMRI, we divided the data between runs, using two runs for the training and the remaining run as the test set. The classifications were repeated three times, using each run as a test set. Thus, there were 32+32 (16 identity-morphs × 2 runs) or 14+14 (7 expression-morphs x 2 runs) exemplars in training sets and 16+16 or 7+7 exemplars in the test sets for each expression and identity analyses, respectively. We employed a different decoding analysis design in fMRI (leave-one-run-out) compared to EEG (leave-one-exemplar-out), because the fMRI data were modelled with a GLM, which might result in dependencies between training and test sets if they contained inputs from same runs. Leave-one-run-out is a standard procedure in fMRI decoding analyses. All decoding analyses were performed separately within each subject.

Decoding analyses of the fMRI data were conducted with The Decoding Toolbox (TDT; Hebart, Görgen, & Haynes, 2015), using beta-images from the first-level GLM in a normalized MNI head space. We used searchlight-based (Kriegeskorte, Goebel, & Bandettini, 2006) decoding with a radius of 10 mm (isotropic), and with default settings of TDT; L2-norm SVM with regularizing parameter C = 1 running in LIBSVM (Chang & Lin, 2011). For EEG data, we used DDTBOX (Bode, Feuerriegel, Bennett, & Alday, 2018) to decode ERPs (event related potentials) averaged over 6 repeated trials (Grootswagers, Wardle, & Carlson, 2017; Isik, Meyers, Leibo, & Poggio, 2014). We used spatiotemporal decoding with 2 consecutive time points, resulting in non-overlapping windows of 10 ms and all 64 channels, resulting in 2*64 = 128 features. Otherwise, default settings of DDTBOX were used; L2-norm SVM with C = 1 running in LIBSVM, same as in fMRI. All decoding analyses were run in Matlab 2018a. After decoding analyses, the results within each pairwise classification were smoothed with FWHM of 3 voxels in fMRI, and with 3 time window (30 ms) moving average in EEG.

### 2.8 Representational similarity analysis

Representational similarity analysis was used to compare the information representations of expressions in fMRI, EEG and models (Figure 1C). A representational dissimilarity matrix (RDM) was calculated for each searchlight voxel in fMRI and for each time window in EEG. The RDMs were calculated separately for each subject, except in the EEG-fMRI-analysis, where the fMRI (but not the EEG) RDMs were averaged over subjects. Each cell *ij* in the RDMs was the decoding accuracy between the two stimulus types of row *i* and column *j*. Three model RDMs were created as follows. First, the *Low-level Model* RDM was built based on filtering all face stimuli with a bank of Gabor filters. Each stimulus image was filtered with 36 filters, using 6 different spatial scales at 6 different orientations. The centre spatial frequency of the filter was varied from 4 to 24 cycles/face width. The spatial frequency bandwidth of the filters was one octave and the orientation bandwidth was 30 degrees. To construct the RDM for expressions, the filter outputs for different identity-morphs were first averaged within each expression-morph, and then the average outputs were correlated with each other and subtracted from one, resulting in the 7×7 expression dissimilarity matrix. In the second model (*Intensity Model*), no differences between different expression categories were assumed; only differences between neutral, half morphs and expressions with full intensities were modelled, with dissimilarity values between neutral and full expressions being 1, between half morphs and neutral/full expressions 0.5, and within each category, 0. Third, the *Category Model* represented categorical processing of emotions, with the 50% morphed and full expressions assumed to be similar, with dissimilarity values between all expressions (irrespective of morph level) being 1, and within expressions 0. The three models were not orthogonal, with Spearman correlations of low-level model with intensity and category models being −.13 and .36, respectively, and with intensity and category model −.11.

The lower triangles of these RDMs were then correlated (Spearman) between fMRI and EEG, fMRI and models, as well as between EEG and models. This was done separately for each fMRI searchlight voxel and each EEG time window and separately for each subject, except for the fMRI-EEG correlations where the subject-averaged fMRI-RDM was used to remove noise and to increase power. We averaged fMRI data instead of EEG as our single-modal EEG results were more robust; similar averaging of MEG data has been used in a previous MEG-fMRI study (Cichy et al., 2016). Mostly similar EEG-fMRI results were obtained when EEG was averaged instead of fMRI (Supplementary Figure 1).

### 2.9 Regions of interest

To further compare and visualize the fMRI-EEG and fMRI-model –correlations, we performed Region of Interest (ROI)-analyses for eight selected areas from the BALSA parcellation map (Glasser et al., 2016): V1, fusiform (face) area (FFC in BALSA), lateral occipital (face) area (LO1 in BALSA), and posterior superior temporal sulcus (pSTS; TPOJ2 in BALSA), all from both hemispheres. In BALSA, the areas are defined multimodally, based on both their anatomical and functional characteristics. The areas were selected based on their known role in face processing (Duchaine & Yovel, 2015), and the results in the ‘Overall information’ –analyses of average face expression processing (see below). From each ROI, five voxels with the highest overall expression information (averaged over subjects, see *2.10. Statistical analysis*) across all 21 pairwise comparisons were selected, and the RDMs from these voxels were averaged for analysis. Thus, the selected ROIs represent the best information available in a given area. As searchlight analysis was used in pairwise classifications, these five voxels contain information from the surrounding voxels as well. While this selection would be circular if analyzing the overall *amount* of information, we used it only to compare the information *structures* (RDMs) in the ROIs to those of models and EEG, and the differences between hemispheres. Supplementary analysis using all the voxels within given BALSA-areas showed similar results (Supplementary Figures 3 & 4A).

### 2.10 Statistical analyses

To calculate a measure for the overall amount of expression information we took the within-subject averages of all the pairwise expression-morph classifications. Similarly, for overall amount of identity information we took the within-subject averages of all the pairwise identity-morph classifications. These (‘Overall information’) averages were calculated in each time window in EEG, and in each searchlight voxel in fMRI. To compare different expressions (‘Expression analyses’), the pairwise classifications of 100% expression (happy, fearful and angry) versus neutral faces were analysed in EEG and in fMRI. The latency differences between different expressions were compared by taking the peak classification accuracies from each subject between 50 – 250 ms and comparing their timings with the Wilcoxon’s signed rank test. For the EEG-fMRI, model-EEG, and model-fMRI analyses, RDMs from each searchlight voxel in fMRI, each time window in EEG, and models, were correlated (EEG-fMRI) or partial-correlated (model-analyses, controlling for other models) using Spearman correlation. For the ROI analyses, exactly same analyses were performed, but instead of all voxels, using the average of 5 voxels from the selected areas. Hemispheric differences were tested with Wilcoxon’s signed rank test, comparing the overall expression information as well as EEG-fMRI –correlations (between 100 – 500 ms) in the ROIs. All statistical significance thresholds were defined by (cluster-based) sign-shuffling permutation tests. In the tests, the decoding accuracies (minus chance level of 50%) or (partial) correlations from each subject were multiplied randomly either by 1 or −1. This was repeated 5000 times, and from each permutation the maximum (cluster) statistic was taken to create a null distribution, FWE-corrected for multiple testing (Nichols & Holmes, 2001). One-sided p<.05 significance threshold was defined as being the top 5% values of the distribution. In EEG, cluster-defining time window threshold was p<.05 (one-sided), and the cluster statistic used was the sum of t-values within each temporal cluster (Maris & Oostenveld, 2007). In fMRI, we used cluster-defining voxel threshold of pseudo-t>3.0, and the cluster statistic used was cluster size (number of voxels). Furthermore, we used variance smoothing of 6 mm FWHM as suggested by Nichols and Holmes (2001). In EEG-fMRI –analyses, above defined cluster analysis was conducted within each time window separately, with no correction applied over the multiple time windows. In ROI-model correlations, the statistic used was partial correlations, and no correction for multiple ROIs were applied. All fMRI statistical analyses were conducted in Matlab, and the permutation analyses were run using SnPM-toolbox for SPM.

### 2.11 Data and code availability

Data and code are available upon request.

## 3 Results

### 3.1 Overall information related to facial expressions and identities

To investigate the processing of facial expression and identities, we showed participants a total of 112 face images in which both emotional expression (neutral, 50% and 100% happy, fearful and angry) and facial identity (4 original identities, 12 morphed identities (33/67%)) varied parametrically (Figure 1A). In order to reveal information related to expressions and identities, we used support vector machine (SVM) decoding to produce pairwise classification accuracies between all stimulus pairs. This was done separately for each searchlight voxel in fMRI with searchlight analysis (Kriegeskorte et al., 2006), and for each time window in EEG (Figure 1C).

To calculate a measure of overall expression processing, we took the mean of all pairwise expression-morph decoding accuracies. In fMRI, this resulted in a large cluster containing the early visual cortex (EVC), occipital face area (OFA), right fusiform face area (FFA) and right posterior superior temporal sulcus (pSTS; Figure 2A). This is in accordance with earlier studies that have shown facial expression processing to be right-lateralized, and occur in the STS, FFA, OFA and EVC (for a review, see Duchaine & Yovel, 2015). The EEG results revealed statistically significant coding of expressions to start at ~120 ms after stimulus onset, and to continue until ~750 ms (Figure 2C) with the highest peak at 270 ms (mean decoding accuracy 54.8%). Previous studies using decoding in MEG, (Dima et al., 2018) and single-cell recordings in monkeys (Sugase et al., 1999) have also found face expression coding to start at ~100 ms.

**Figure 2.**
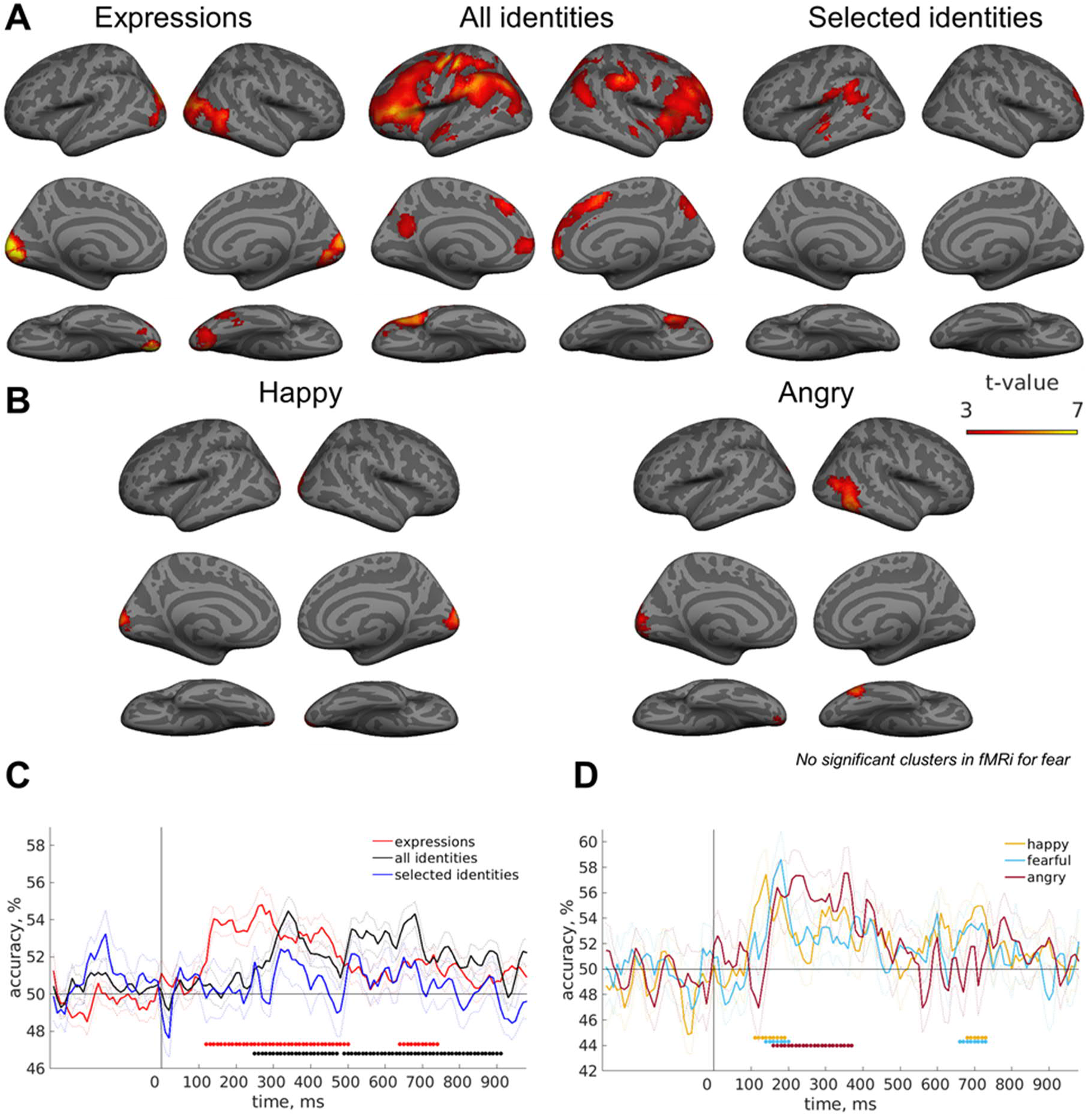
Means of pairwise decoding accuracies of expressions and identities. **A)** In fMRI, a large cluster for expression classifications was found in bilateral occipital cortex and right temporal lobe, including OFA, FFA and pSTS. Identity results were possibly obscured by the task and motor responses (middle column, ‘All identities’). Identities with similar responses (rightmost column, ‘Selected identities’) revealed no clusters in expected face regions. **B)** When different expressions were analyzed separately, we found significant coding of happy (vs. neutral) expressions in EVC, and angry faces in EVC, right pSTS and right FFA. No significant clusters for fear were found. fMRI cluster-forming threshold pseudo-t>3 (variance-smoothed), cluster-level p(FWE)<.05. **C)** Average decoding accuracies in EEG. **D)** In EEG, the decoding accuracies for different expressions peaked between 130 – 200 ms after stimulus onset. Dots in EEG mark significant time windows (cluster weight analysis, cluster-forming threshold p<.05, one-sided; FWE-corrected), and dotted lines depict SEMs.

For face identities, we performed similar EEG and fMRI analyses. Our analyses suggested that the motor responses of the task of gender identification were obscuring the identity results, showing for example significant decoding accuracies in fMRI in the left motor cortex (Figure 2A, ‘All identities’). To account for this, we performed analyses using only the within-gender identity-morph pairs in which participants answered correctly (using the same finger) over 90% of the trials (‘Selected identities’). Only 17/120 comparison pairs fulfilled the criteria. The results showed significant clusters in the left temporo-parietal junction and anterior STS, but not in the visual areas or in the ventral temporal areas implicated in face-specific processing (Figure 2A), or in any time window in EEG. Thus, as no expected face patches showed significant decoding results, no further analyses for facial identities were performed.

### 3.2 Different temporal and spatial responses to different facial expressions

To compare different expressions, we examined the classifications of neutral expression compared separately to (100%) happy, fearful, and angry faces. Happy faces were significantly classified from early visual cortex in fMRI (Figure 2B), with the highest peak in EEG at 140 ms (57.5%; Figure 2D). Angry faces peaked later in EEG, at 220 ms and 350 ms (57.4% and 57.5%, respectively; Figure 2D), and were accurately classified in fMRI both from V1 and from a more anterior cluster near the right pSTS and FFA (Figure 2B). Accurate classification of fearful faces in the EEG data was found in time windows between those found for happy and angry faces, with the highest peak at 180 ms (58.6%), while no significant clusters were found with fMRI. To statistically test temporal differences in the three expressions, we compared the latencies of the highest decoding accuracies of each expression. Decoding accuracy of happy faces peaked significantly earlier than both fearful (p=.021) and angry (p=.005) faces, while no statistically significant difference was found between fearful and angry expressions (p=.36). Furthermore, while happy and fearful faces were significantly classified only in a relatively short (<100 ms) time window, the classification of angry faces remained significant for 220 ms (Figure 2D).

### 3.3 Combined EEG-fMRI reveals spatiotemporal pattern of face processing

We used representational similarity analysis (Kriegeskorte et al., 2008) to compare the information structures related to the processing of facial expressions in fMRI and EEG. We correlated representational dissimilarity matrixes (RDMs) from each searchlight voxel in fMRI and each time window in EEG. Each cell of the RDMs was the decoding accuracy between two stimuli in a certain searchlight voxel or in a certain time window. We found that information in EEG correlated significantly with activity patterns in early visual cortex from 130 ms onwards (Figure 3). At 190 – 250 ms, significant correlations were found in parietal areas, the EVC as well as in left FFA, OFA and part of left STS. These results are consistent with temporal progress of information processing from posterior to more anterior sites in visual object categorization (Cichy et al., 2016).

**Figure 3.**
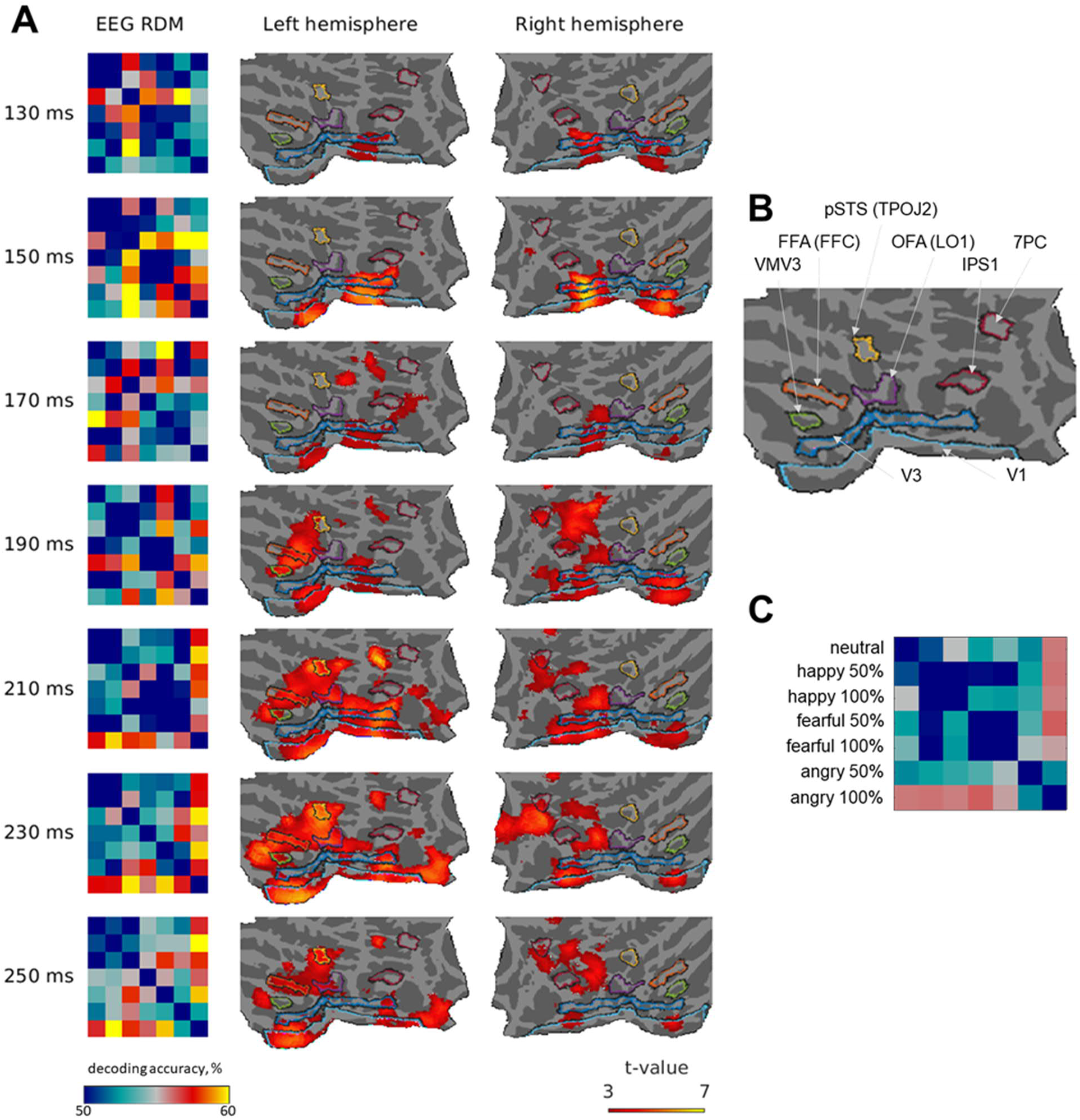
EEG-fMRI RDM correlations. **A)** In each time window, from left to right: EEG RDM, correlation between EEG and fMRI RDMs in each voxel in left and right hemisphere. The fMRI RDMs are the same in each time window. The flatmaps contain approximately the posterior half of the brain. Cluster analysis with cluster-forming threshold pseudo-t>3 (variance-smoothed), cluster-level p(FWE)<.05, each time window analysed separately. No significant clusters were found before 130 ms, and only sparsely after 250 ms. **B)** Labels for marked flatmap brain regions (from BALSA parcellation). **C)** Labels for RDM rows/columns.

To further compare the information structures between fMRI and EEG, we calculated RDMs (Figure 4A) for the mean of five searchlight voxels having highest overall expression information in different ROIs (Figure 4B), and correlated them to the EEG data. Similarly to the whole brain searchlight analysis, V1 had the highest correlation with the earliest time windows (~120 – 250 ms), peaking around 150 ms (Figure 4C). Bilateral FFA, OFA and left pSTS all peaked later, around 240 ms. In the right pSTS, no significant correlations were found. In left OFA, the correlations remained significant until 480 ms. The overall spatio-temporal structure found here was quite similar to that reported earlier in a MEG study (Vida et al., 2017) looking at the coding of face identities in the brain, finding accurate classifications in the left V1 around 150 ms, and relatively higher decoding accuracy for the right LO and the right FG around 250 ms. To ensure that our results were not due to selecting the five most informative voxels, the ROI analyses were repeated using all the voxels within a given BALSA-area. The results were similar except that the right FFA did not correlate significantly with EEG (Supplementary Figure 3). Finally, as the whole brain EEG-fMRI –results (Figure 3) seemed to show a hemispheric bias to the left, we tested hemispheric differences in the ROIs. We found higher correlations with EEG in the left than right hemisphere in OFA (p=.031) and pSTS (p=.006), but not in FFA (p=.055) or V1 (p=.65), and, when taking all the voxels within the ROIs, for OFA (p=.040), FFA (p=.010) and pSTS (p=.004). However, comparing the overall expression information in the ROIs showed no left bias, with no differences between hemispheres in V1, OFA, or FFA (all ps>.08), and higher average decoding accuracies in pSTS in the right than left hemisphere (p=.017).

**Figure 4.**
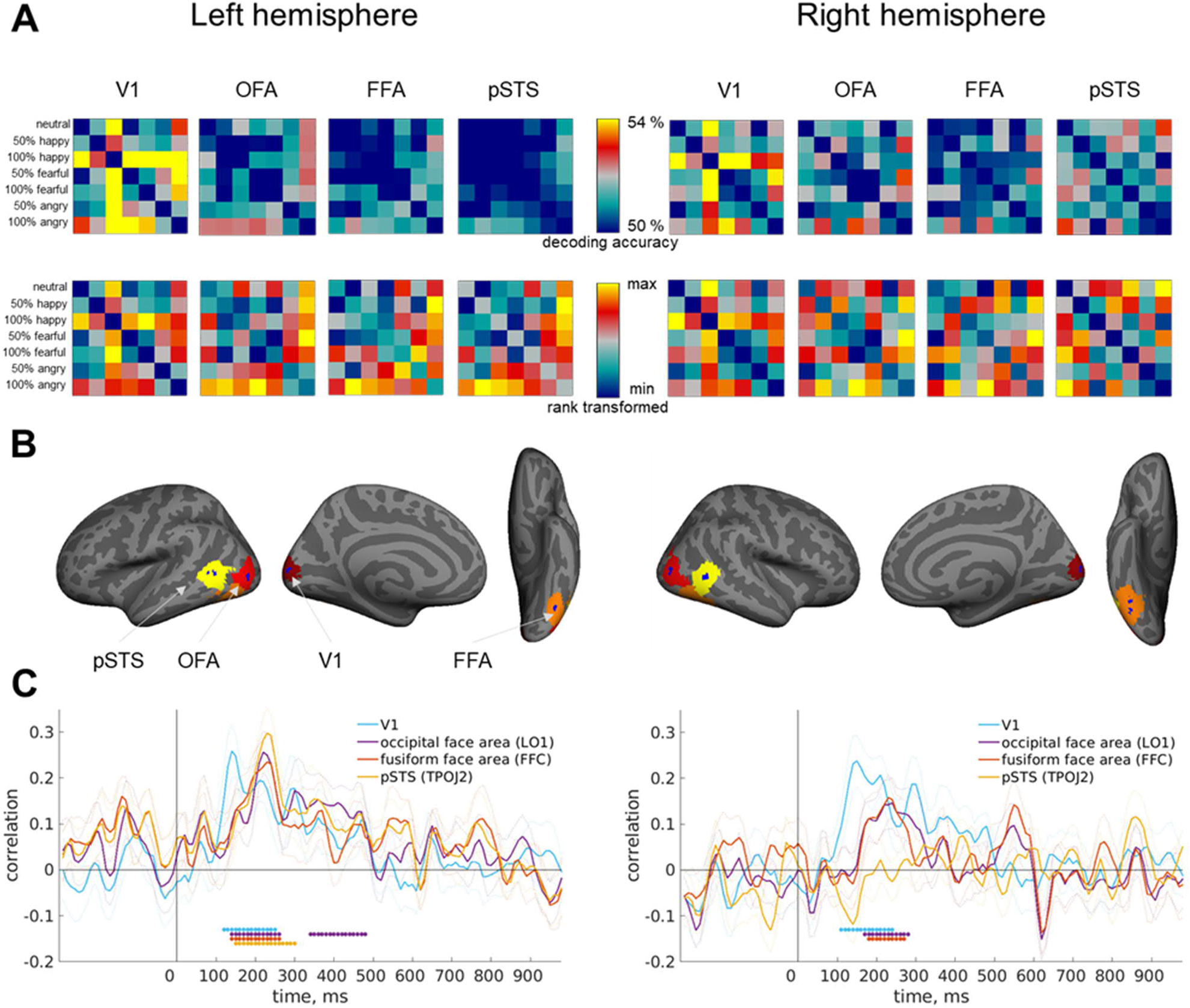
ROI-analysis. **A)** RDMs from selected ROIs. In the top row, decoding accuracies. In the bottom row, rank-transformed RDMs (sorted according to their rank from lowest to highest) showing the relative weight of each decoding pair within each ROI. **B)** Locations of ROIs. Within every ROI (BALSA-area), 5 voxels that had the highest overall expression information over all decoding pairs were selected. Voxel locations shown in dark blue and the total area of their searchlight in yellow, orange and red. **C)** Correlations between each fMRI ROI and EEG from both hemispheres. Dots mark significant time windows (cluster weight analysis, cluster-forming threshold p<.05, one-sided; FWE-corrected), and dotted lines depict SEMs.

### 3.4 Models for expression processing

We compared our data with the three model RDMs (Figure 5A). Similarly to the EEG-fMRI –correlations, we correlated each searchlight voxel from fMRI (Figure 5B) and each time window from EEG (Figure 5C) with each of the models, using partial correlations controlling for the other models. The *Low-level Model* correlated with EVC in fMRI (Figure 5, top row), and with EEG between 110 – 290 ms. The *Intensity Model*, on the other hand, had the most pronounced correlation with an area near the right pSTS (Figure 5, middle row). In EEG, the intensity model did not correlate significantly with information in any time window (Figure 5C). This is consistent with the EEG-fMRI-analysis also finding no correlations with right STS and EEG. The *Category Model* did not correlate significantly with fMRI nor EEG data. Our results suggest that the classifications in EEG were mostly based on information about low-level visual features, and in fMRI the classifications were based on low-level features as well as expression intensities. As several studies have found the right pSTS to be especially sensitive to differences between neutral and emotional faces (Greening et al., 2018; Zhang et al., 2016), we calculated the model correlations with fMRI data without the neutral faces (Supplementary Figures 2 and 4B). The results were similar, showing that the model correlations were not driven only by separating neutral faces from emotional faces, but instead by the intensity of the expressions.

**Figure 5.**
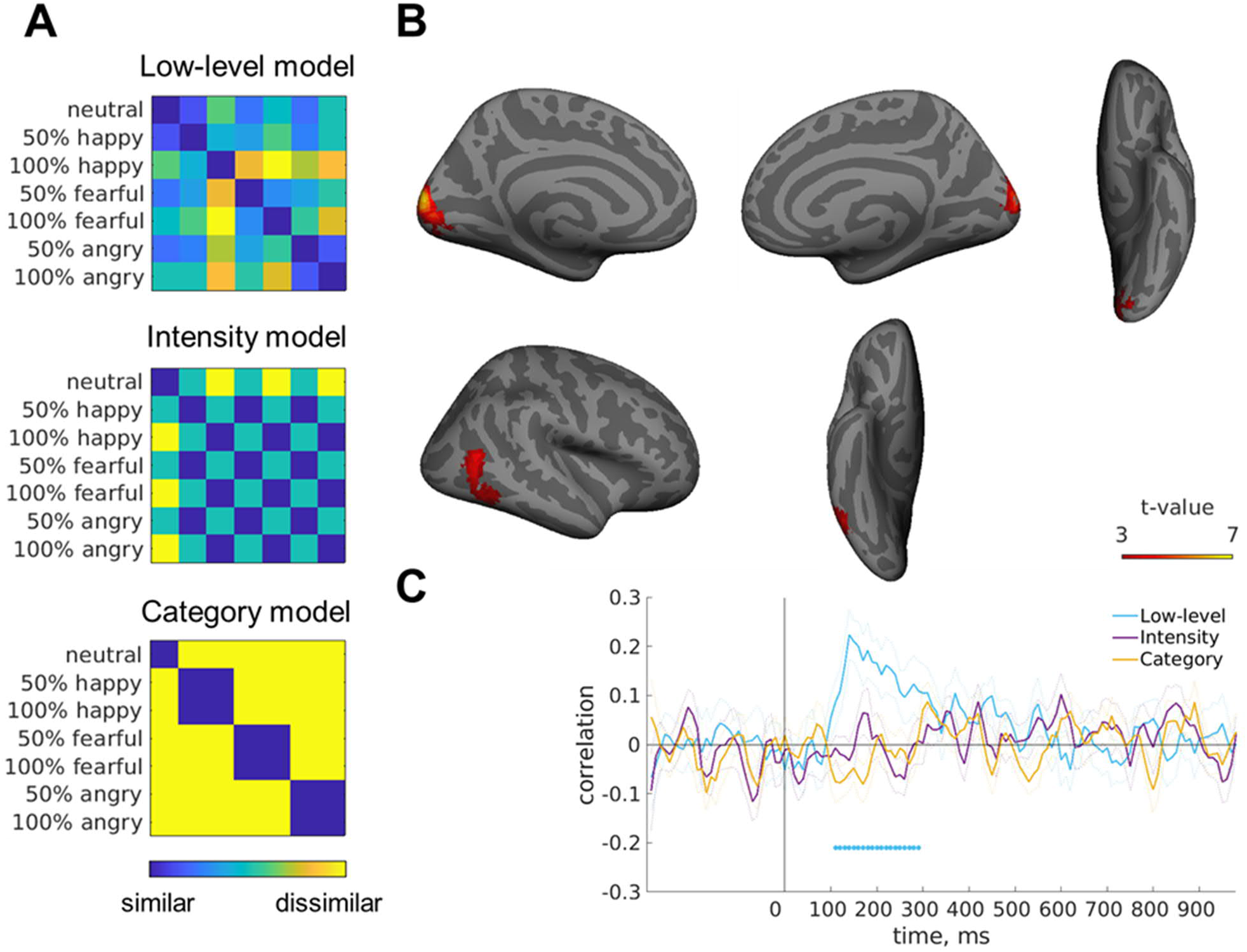
Model analysis. **A)** RDMs for the three models. **B)** fMRI partial correlations with the models (no significant clusters for category model were found). Cluster analysis with cluster-forming threshold pseudo-t>3 (variance-smoothed), cluster-level p(FWE)<.05. **C)** Partial correlations between EEG and the models. Dots mark significant time windows (cluster weight analysis, cluster-forming threshold p<.05, one-sided; FWE-corrected), and dotted lines depict SEMs.

Finally, we compared the models with the searchlight voxels with maximal information in each ROI. As seen in Figure 6, these results were mainly similar to the results of the whole brain correlations. The low-level model correlated with V1 and the intensity model with the right pSTS. While these results were consistent with the results of the whole brain model-searchlight, they also revealed the coding of emotion intensity in the right FFA, emotion category in the right pSTS and left OFA, and low-level features in the left FFA, not apparent in the whole brain results. We conducted three control analyses. First, we tested model correlations when taking all searchlight voxels (instead of the five most informative) from a given BALSA area. The results were identical (Supplementary Figure 4A). Second, we repeated the analyses without neutral faces and the results were again highly similar (Supplementary Figure 4B). Finally, in our *Category Model*, the 50% morphed expressions were categorized with the full expressions as they, subjectively assessed, looked more like an expressive than a neutral face (Figure 1B). When they were instead categorized to be similar with the neutral faces, the EEG and whole brain results did not change, but there were some minor changes in the ROI results (Supplementary Figure 4C).

**Figure 6.**
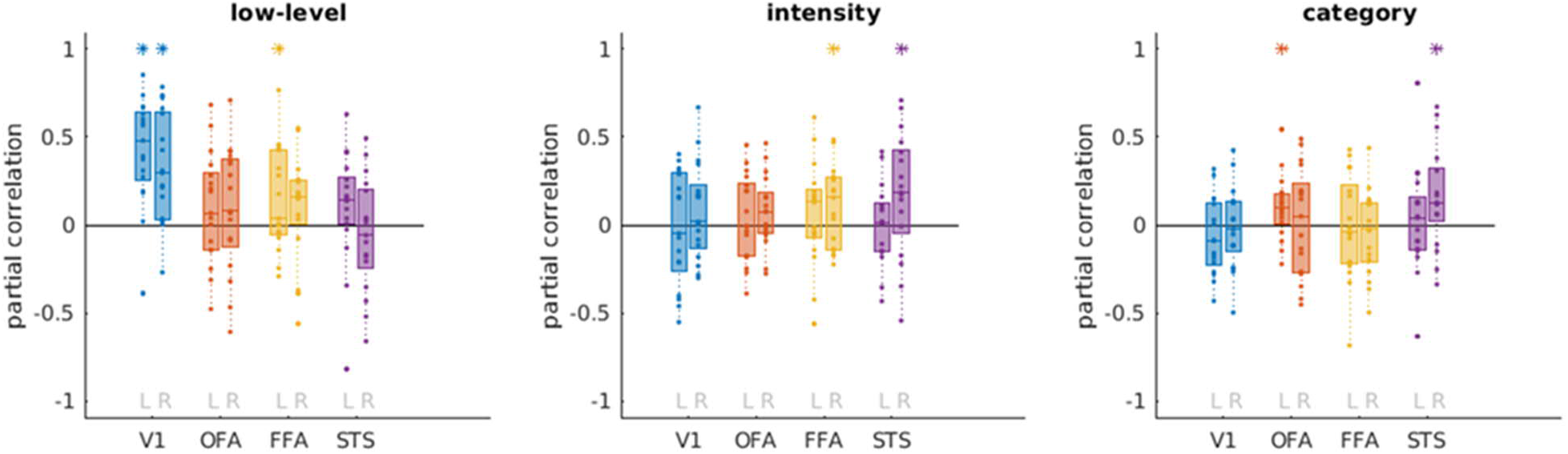
Model-ROI partial correlations. Boxes indicate data from 25^th^ to 75^th^ percentiles, lines the medians, dots individual subjects, and asterisks mark significance (sign-shuffled p<.05, one-sided). L and R refer to the left and right hemisphere, respectively, OFA to occipital face area (LO1 in BALSA), FFA to fusiform face area (FFC in BALSA) and STS to posterior superior temporal sulcus (TPOJ2 in BALSA).

## 4 Discussion

We investigated the processing of facial expressions and identities in the brain with EEG and fMRI. We used SVM pairwise decoding to measure the amount of information between each stimulus, and compared the information patterns in fMRI, EEG, and models by correlating dissimilarity matrices obtained with the different methods. Our results from EEG-fMRI-correlations showed spatio-temporal spreading of expression information from EVC at around 130 ms to left FFA, OFA and pSTS, which correlated with the EEG data from 190 ms onwards. Comparisons between models and the EEG data suggest that EEG contains more information about low-level visual features than expression intensities or categories. Comparisons between models and the fMRI data, however, indicate low-level visual feature-based processing in the early visual cortex and expression intensity-based processing in the right pSTS. Thus, our results suggest gradual change in the processing of facial information from low-level visual features in the early visual cortex to facial feature analysis in the left inferior temporal areas and expression intensity analysis in the right pSTS.

For identity information, our results were likely compromised by the motor task of responding to the gender of the shown face. However, as we used searchlight analyses in the fMRI data, limiting the information used in each classification to a small set of voxels, the motor activations from the parietal and frontal lobes should not dilute possible findings in the visual and face processing areas most important for the aim of our study. Nevertheless, no significant classification of identity information was found from these areas, possibly due to the smaller variance of visual information between the identities than the expressions. In EEG, although we found processing of identities from 250 ms onwards, these results vanished when we tried to control motor responses by selecting only a subset of the identity pairwise classifications. Thus, no further analyses or conclusions relating to the identity coding were made. It is worth noting that our decoding design required some invariance to expressions in the identity classifications. Some previous studies with invariance requirement to viewpoints (Guntupalli et al., 2017) and generalization across face halves (Anzellotti & Caramazza, 2016), have found identity coding in the right inferior frontal and anterior temporal lobe, respectively, but not in FFA or OFA (although see Anzellotti, Fairhall, & Caramazza, 2014 for rotation-invariant classification in OFA and FFA). However, as the area in the frontal cortex is harder to disentangle from the motor responses, and our fMRI signal was weak in the anterior temporal lobe, our results cannot weigh in on the role of these areas in identity processing.

Analyzing our expression results more closely, we first looked at the areas and timing of overall expression information. In EEG, expressions were decoded significantly between ~120 – 750 ms, replicating earlier findings of expression processing starting roughly around 100 ms (Dima et al., 2018; Müller-Bardorff et al., 2018; Sugase et al., 1999). While several studies do not report such an early effect of expression processing (e.g., Blau, Maurer, Tottenham, & Mccandliss, 2007; Müller-Bardorff et al., 2016), this can be explained with the less sensitive methods mainly used in those studies. In fMRI, significant classification was found, as expected, from early visual areas and the face processing areas in the right fusiform gyrus (FFA), lateral occipital cortex (OFA) and near posterior superior temporal sulcus (pSTS). For the separate expressions, happy faces were accurately classified in early visual areas and most pronouncedly at early time windows (~140 ms), while information on angry faces was found, in addition to EVC, from a cluster near the right pSTS and FFA, and mainly later in time (~200 – 350 ms). Highest classification accuracy for fearful faces was found in EEG in a later time window than for happy faces, but earlier than for angry faces (~180 ms), and no significant clusters were found in fMRI. In addition to the differences in peak latencies, the coding of angry faces remained statistically significant longer than other expressions, possibly showing the evolutionary importance of paying attention to angry faces, longer-lasting processing of eye than mouth information, or both.

We used representational similarity analysis to compare the information structure in EEG and in fMRI. One of our main findings was a spreading of information similarity from early visual cortex and early in time (120 – 150 ms) to more anterior areas in left (ventral) temporal lobe and later in time (190 – 250 ms). Face models (Duchaine & Yovel, 2015; Haxby et al., 2000) postulate spatio-temporal progress during the processing of faces. However, as every non-invasive method (fMRI, M/EEG) has a rather weak spatial or temporal resolution, few studies have been able to focus on both dimensions at once. Thus, while there is an exhaustive literature on both spatial areas of face processing from fMRI studies, and temporal components found using M/EEG, the linkage between these results is much less understood. Our study takes a step into that direction, being the first to our knowledge to combine M/EEG and fMRI with RSA to understand the processing of facial expressions. Interestingly, we found EEG-fMRI correlations mainly in the left hemisphere, which is less involved in face processing. We hypothesize that this might be due to our EEG data reflecting more lower level visual features, as shown in our model analyses. Our model results suggest that in the right hemisphere, at least in the right pSTS, there is higher-level representations of face expressions. As these were not found from the EEG data, that could explain the lower correlations between EEG and fMRI in the right than the left hemisphere. In ROI analyses we similarly found an early peak in fMRI-EEG correlations in V1, and a later peak in OFA and FFA. Our time trajectories are similar to those reported in a previous MEG study (Vida et al., 2017), which found first peaks for V1, OFA and FFA at around 150 ms and later peaks ~250 ms. Both in our and their results, information in V1 clearly dominated early in time, and later in time, the information from OFA and FFA was relatively more prominent, though mainly not surpassing correlations with V1.

We compared our data to three models depicting low-level visual information, emotional intensity, and emotion category information respectively, while controlling for other models by using partial correlation. The low-level model matched with the information in the early visual cortex and EEG data from ~140 ms to 290 ms. The intensity model, on the other hand, correlated with the right pSTS and not at all with EEG. The category model did not correlate with either EEG or fMRI data. As we found no EEG-fMRI correlations from the right pSTS, these model correlations provide an explanation for that: the right pSTS seemed to code emotional intensity with less sensitivity than the lower level features, and this coding was not separable in EEG data. Previous studies have shown STS to be especially sensitive to differences between neutral and emotional faces (Greening et al., 2018; Zhang et al., 2016). Our results suggest that this is due to STS coding expression intensity, and thus provides a more specific model for the role of pSTS in face processing. Srinivasan, Golomb, and Martinez (2016) found pSTS to be especially sensitive to action units in the face. As we did not model action units, future studies should disentangle whether the intensity and action unit coding reflect the same or different processes in pSTS.

It is worth emphasizing what the RDM correlations between EEG and fMRI measure exactly, and what the underlying assumptions are. In the EEG-fMRI analysis, whole brain EEG RDMs (that may contain multiple, spatially different sources) are compared to local fMRI RDMs (that contain information from a temporally long period), and this might cause spurious correlations. However, our (as well as those of others) EEG-fMRI –correlation results are surprisingly systematic, and follow, for example, the known processing hierarchy in visual cortex. Still, the EEG-fMRI correlations should be interpreted with caution. The RDM correlations are insensitive to the *amount* of information, and measure only the information structure, namely the *relative* information between the different expressions. This is contrary to the single-modality classifications, which show the amount of information, but not the relative information. Thus, it is entirely possible for there to be an area in the brain having high decoding accuracy for all of the different expressions, for example, but still not showing up in the model or in the EEG-fMRI correlations. One example of these inconsistencies in our data is the higher EEG-fMRI correlations in the *left* compared to the right hemisphere, while the amount of overall expression information differed only in pSTS, with more information in the *right* hemisphere. Thus, the hemispheric differences in the EEG-fMRI –correlations show only that the left hemisphere had a more similar information structure to the EEG data than the right hemisphere, and not that it was more involved in the processing of faces. Furthermore, our model analyses suggest that the EEG data contained mostly information related to low-level visual features, and therefore the EEG-fMRI correlations likely mostly reflect visual feature analyses related to the processing of facial expression. These examples also show how different methods (EEG, fMRI, models) complement each other.

In conclusion, we showed a glimpse of the spatio-temporal coding of face expressions in the brain. Starting from the early visual cortex, the processing spreads to OFA, FFA and pSTS around 200 ms. By comparing these to the unimodal results from EEG and fMRI, as well as model correlations, our results suggest that the processing of faces hierarchically changes from visual features in early visual cortex to expression intensities in the right pSTS. As slightly different results were obtained with EEG/fMRI classifications and EEG-fMRI/model correlations, our results highlight the importance of combining M/EEG and fMRI data and models in order to understand the spatio-temporal processing dynamics in the human brain.

## Acknowledgements

This work was supported by an Academy of Finland grant to VRS (grant number 298329).

## References

Ambrus, G. G., Kaiser, D., Cichy, R. M., & Kovács, G. (2019). The Neural Dynamics of Familiar Face Recognition. Cerebral Cortex. https://doi.org/10.1093/cercor/bhz010

Anzellotti, S., & Caramazza, A. (2016). From Parts to Identity: Invariance and Sensitivity of Face Representations to Different Face Halves. Cerebral Cortex, 26(5), 1900–1909. http://doi.org/10.1093/cercor/bhu337

Anzellotti, S., Fairhall, S. L., & Caramazza, A. (2014). Decoding representations of face identity that are tolerant to rotation. Cerebral Cortex, 24(8), 1988–1995. http://doi.org/10.1093/cercor/bht046

Atkinson, A. P., & Adolphs, R. (2011). The neuropsychology of face perception: beyond simple dissociations and functional selectivity. Philosophical Transactions of the Royal Society of London. Series B, 366(1571), 1726–1738. http://doi.org/10.1098/rstb.2010.0349CerebralCortex, 24(8), 1988–1995.

Bernstein, M., & Yovel, G. (2015). Two neural pathways of face processing: A critical evaluation of current models. Neuroscience and Biobehavioral Reviews, 55, 536–546. http://doi.org/10.1016/j.neubiorev.2015.06.010

Blau, V. C., Maurer, U., Tottenham, N., & Mccandliss, B. D. (2007). The face-specific N170 component is modulated by emotional facial expression. Behavioral and Brain Functions, 13, 1–13. http://doi.org/10.1186/1744-9081-3-7

Bode, S., Feuerriegel, D., Bennett, D., & Alday, P. M. (2018). The Decision Decoding ToolBOX (DDTBOX) – A Multivariate Pattern Analysis Toolbox for Event-Related Potentials. Neuroinformatics, 1–16. http://doi.org/10.1007/s12021-018-9375-z

Bushnell, I. W. R. (2001). Mother’s Face Recognition in Newborn Infants: Learning and Memory. Infant and Child Development, 10, 67–74. http://doi.org/10.1002/icd.248

Bushnell, I. W. R., Sai, F., & Mullin, J. T. (1989). Neonatal recognition of the mother’s face. British Journal of Developmental Psychology, 7(1), 3–15. http://doi.org/10.1111/j.2044-835X.1989.tb00784.x

Carlin, J. D., & Kriegeskorte, N. (2017). Adjudicating between face-coding models with individual-face fMRI responses. PLoS Computational Biology, 13(7), 1–28. http://doi.org/10.1371/journal.pcbi.1005604

Chang, C.-C., & Lin, C.-J. (2011). LIBSVM: A Library for Support Vector Machines. ACM Transactions on Intelligent Systems and Technology, 2(3), 1–27. http://doi.org/10.1145/1961189.1961199

Chang, L., & Tsao, D. Y. (2017). The Code for Facial Identity in the Primate. Cell, 169(6), 1013–1020.e14. http://doi.org/10.1016/j.cell.2017.05.011

Cichy, R. M., & Pantazis, D. (2017). Multivariate pattern analysis of MEG and EEG: A comparison of representational structure in time and space. NeuroImage, 158, 441–454. http://doi.org/10.1016/J.NEUROIMAGE.2017.07.023

Cichy, R. M., Pantazis, D., & Oliva, A. (2014). Resolving human object recognition in space and time. Nature Neuroscience, 17(3), 455–62. http://dx.doi.org/10.1038/nn.3635

Cichy, R. M., Pantazis, D., & Oliva, A. (2016). Similarity-Based Fusion of MEG and fMRI Reveals Spatio-Temporal Dynamics in Human Cortex During Visual Object Recognition. Cerebral Cortex, 26(8), 3563–3579. http://doi.org/10.1093/cercor/bhw135

Collins, J. A., & Olson, I. R. (2015). Beyond the FFA: The Role of the Ventral Anterior Temporal Lobes in Face Processing. Neuropsychologia, 65–79. http://doi.org/10.1016/j.neuropsychologia.2014.06.005.Beyond

Cortes, C., & Vapnik, V. (1995). Support-Vector Networks. Machine Learning, 20(3), 273–297. http://doi.org/10.1007/BF00994018

Dima, D. C., Perry, G., Messaritaki, E., Zhang, J., & Singh, K. D. (2018). Spatiotemporal dynamics in human visual cortex rapidly encode the emotional content of faces. Human Brain Mapping. http://doi.org/10.1002/hbm.24226

Dobs, K., Schultz, J., Bülthoff, I., & Gardner, J. L. (2018). Task-dependent enhancement of facial expression and identity representations in human cortex. NeuroImage, 172(July 2017), 689–702. http://doi.org/10.1016/j.neuroimage.2018.02.013

Dobson, V., & Teller, D. Y. (1978). Visual acuity in human infants: A review and comparison of behavioral and electrophysiological studies. Vision Research, 18(11), 1469–1483. http://doi.org/10.1016/0042-6989(78)90001-9

Duchaine, B., & Yovel, G. (2015). A Revised Neural Framework for Face Processing. Annual Review of Vision Science, 1(1), 393–416. http://doi.org/10.1146/annurev-vision-082114-035518

Ebner, N. C., Riediger, M., & Lindenberger, U. (2010). FACES - A database of facial expressions in young, middle-aged, and older women and men: Development and validation. Behavior Research Methods, 42(1), 351–62. http://doi.org/10.3758/BRM.42.1.351

Edelman, S., Grill-Spector, K., Kushnir, T., & Malach, R. (1998). Toward direct visualization of the internal shape represetation space by fMRI. Psychobiology, 26(4), 309–321. https://doi.org/10.1093/cercor/11.10.946

Freiwald, W. A., & Tsao, D. Y. (2010). Functional Compartmentalization and Viewpoint Generalization Within the Macaque Face-Processing System. Science, 330(5), 845–851. https://doi.org/10.1126/science.1229223

Glasser, M. F., Coalson, T. S., Robinson, E. C., Hacker, C. D., Harwell, J., Yacoub, E., … Van Essen, D. C. (2016). A multi-modal parcellation of human cerebral cortex. Nature, 536(7615), 171–178. http://doi.org/10.1038/nature18933

Gold, J. M., Mundy, P. J., & Tjan, B. S. (2012). The Perception of a Face Is No More Than the Sum of Its Parts. Psychological Science, 23(4), 427–434. http://doi.org/10.1177/0956797611427407

Greening, S. G., Mitchell, D. G. V., & Smith, F. W. (2018). Spatially generalizable representations of facial expressions: Decoding across partial face samples. Cortex, 101, 31–43. http://doi.org/10.1016/j.cortex.2017.11.016

Grootswagers, T., Wardle, S. G., & Carlson, T. A. (2017). Decoding Dynamic Brain Patterns from Evoked Responses: A Tutorial on Multivariate Pattern Analysis Applied to Time Series Neuroimaging Data. Journal of Cognitive Neuroscience, 29(4), 677–697. https://doi.org/10.1162/jocn

Guntupalli, J. S., Wheeler, K. G., & Gobbini, M. I. (2017). Disentangling the Representation of Identity from Head View Along the Human Face Processing Pathway. Cerebral Cortex, 27, 46–53. http://doi.org/10.1093/cercor/bhw344

Haxby, J. V., Hoffman, E. A., & Gobbini, M. I. (2000). The distributed human neural system for face perception. Trends in Cognitive Sciences, 4(6), 223–233. http://doi.org/10.1016/S1364-6613(00)01482-0

Hebart, M. N., Bankson, B.B., Harel, A., Baker, C. I., & Cichyl, R. M. (2018). The representational dynamics of task and object processing in humans. eLife, 7, 1–21. https://doi.org/10.7554/eLife.32816

Hebart, M. N., Görgen, K., & Haynes, J.-D. (2015). The Decoding Toolbox (TDT): a versatile software package for multivariate analyses of functional imaging data. Frontiers in Neuroinformatics, 8, 88. http://doi.org/10.3389/fninf.2014.00088

Henriksson, L., Mur, M., & Kriegeskorte, N. (2015). Faciotopy-A face-feature map with face-like topology in the human occipital face area. Cortex, 72, 156–167. http://doi.org/10.1016/j.cortex.2015.06.030

Hinojosa, J. A., Mercado, F., & Carretié, L. (2015). N170 sensitivity to facial expression: A meta-analysis. Neuroscience & Biobehavioral Reviews, 55, 498–509. http://doi.org/10.1016/J.NEUBIOREV.2015.06.002

Isik, L., Meyers, E. M., Leibo, J. Z., & Poggio, T. (2014). The dynamics of invariant object recognition in the human visual system. Journal of Neurophysiology, 111(1), 91–102. https://doi.org/10.1152/jn.00394.2013

Kanwisher, N., & Yovel, G. (2006). The fusiform face area: a cortical region specialized for the perception of faces. Philosophical Transactions of the Royal Society of London. Series B, 361(1476), 2109–28. http://doi.org/10.1098/rstb.2006.1934

Kietzmann, T. C., Spoerer, C. J., Sörensen, L., Cichy, R. M., Hauk, O., & Kriegeskorte, N. (2019). Recurrence required to capture the dynamic computations of the human ventral visual stream. arXiv:1903.05946.

Kothe, C. (2013). The artifact subspace reconstruction method. http://sccn.ucsd.edu/eeglab/plugins/ASR.pdf

Kriegeskorte, N., Formisano, E., Sorger, B., & Goebel, R. (2007). Individual faces elicit distinct response patterns in human anterior temporal cortex. Proceedings of the National Academy of Sciences of the United States of America, 104(51), 20600–20605. https://doi.org/10.1073/pnas.0705654104

Kriegeskorte, N., Goebel, R., & Bandettini, P. a. (2006). Information-based functional brain mapping. Proceedings of the National Academy of Sciences of the United States of America, 103(10), 3863–8. http://doi.org/10.1073/pnas.0600244103

Kriegeskorte, N., Mur, M., & Bandettini, P. a. (2008). Representational similarity analysis - connecting the branches of systems neuroscience. Frontiers in Systems Neuroscience, 2, 4. http://doi.org/10.3389/neuro.06.004.2008

Maris, E., & Oostenveld, R. (2007). Nonparametric statistical testing of EEG-and MEG-data. Journal of Neuroscience Methods, 164(1), 177–190. https://doi.org/10.1016/j.jneumeth.2007.03.024

Mullen, T (2012). CleanLine EEGLAB plugin. San Diego, CA: Neuroimaging Informatics Toolsand Resources Clearinghouse (NITRC).

Müller-Bardorff, M., Bruchmann, M., Mothes-Lasch, M., Zwitserlood, P., Schlossmacher, I., Hofmann, D., Straube, T. (2018). Early brain responses to affective faces: A simultaneous EEG-fMRI study. NeuroImage, 178, 660–667. http://doi.org/10.1016/J.NEUROIMAGE.2018.05.081

Müller-Bardorff, M., Schulz, C., Peterburs, J., Bruchmann, M., Mothes-Lasch, M., Miltner, W., & Straube, T. (2016). Effects of emotional intensity under perceptual load: An event-related potentials (ERPs) study. Biological Psychology, 117, 141–149. http://doi.org/10.1016/j.biopsycho.2016.03.006

Nemrodov, D., Behrmann, M., Niemeier, M., Drobotenko, N., & Nestor, A. (2019). Multimodal evidence on shape and surface information in individual face processing. NeuroImage, 184, 813–825. http://doi.org/10.1101/299933

Nichols, T. E., & Holmes, A. P. (2001). Nonparametric permutation tests for functional neuroimaging: A primer with examples. Human Brain Mapping, 15, 1–25. https://doi.org/10.1002/hbm.1058

Pitcher, D., Walsh, V., & Duchaine, B. (2011). The role of the occipital face area in the cortical face perception network. Experimental Brain Research, 209(4), 481–493. http://doi.org/10.1007/s00221-011-2579-1

Reid, V. M., Dunn, K., Young, R. J., Amu, J., Donovan, T., & Reissland, N. (2017). The Human Fetus Preferentially Engages with Face-like Visual Stimuli. Current Biology, 27(12), 1825–1828. http://doi.org/10.1016/j.cub.2017.05.044

Richler, J. J., & Gauthier, I. (2014). A Meta-Analysis and Review of Holistic Face. Psychological Bulletin, 140(5), 1281–1302. http://doi.org/10.1037/a0037004.A

Rossion, B., & Jacques, C. (2011). The N170: understanding the time-course of face perception in the human brain. The Oxford Handbook of ERP Components, Oxford University Press. 115–142. http://doi.org/10.1093/oxfordhb/9780195374148.013.0064

Sadeh, B., Podlipsky, I., Zhdanov, A., & Yovel, G. (2010). Event-related potential and functional MRI measures of face-selectivity are highly correlated: A simultaneous ERP-fMRI investigation. Human Brain Mapping, 31(10), 1490–1501. http://doi.org/10.1002/hbm.20952

Said, C. P., Haxby, J. V., & Todorov, A. (2011). Brain systems for assessing the affective value of faces. Philosophical Transactions of the Royal Society of London B, 366(1571), 1660–1670. http://doi.org/10.1098/rstb.2010.0351

Said, C. P., Moore, C. D., Engell, A. D., Todorov, A., & Haxby, J. V. (2010). Distributed representations of dynamic facial expressions in the superior temporal sulcus. Journal of Vision, 10(5), 11. https://doi.org/10.1167/10.5.11

Salmela, V., Salo, E., Salmi, J., & Alho, K. (2018). Spatiotemporal Dynamics of Attention Networks Revealed by Representational Similarity Analysis of EEG and fMRI. Cerebral Cortex, 28(2), 549–560. http://doi.org/10.1093/cercor/bhw389

Schweinberger, S. R., & Neumann, M. F. (2016). Repetition effects in human ERPs to faces. Cortex, 80, 141–153. http://doi.org/10.1016/J.CORTEX.2015.11.001

Shen, J., & Palmeri, T. J. (2015). The Perception of a Face is Greater Than the Sum of Its Parts The perception of a face is no more than the sum of its parts. Psychonomic Bulletin & Review, 22, 710–716. http://doi.org/10.3758/s13423-014-0726-y

Srinivasan, R., Golomb, J. D., & Martinez, A. M. (2016). A Neural Basis of Facial Action Recognition in Humans. Journal of Neuroscience, 36(16), 4434–4442. https://doi.org/10.1523/JNEUROSCI.1704-15.2016

Sugase, Y., Yamane, S., Ueno, S., & Kawano, K. (1999). Global and fine information coded by single neurons in the temporal visual cortex. Nature, 400(6747), 869–873. http://doi.org/10.1038/23703

Surguladze, S. A., Brammer, M. J., Young, A. W., Andrew, C., Travis, M. J., Williams, S. C. R., & Phillips, M. L. (2003). A preferential increase in the extrastriate response to signals of danger. NeuroImage, 19(4), 1317–1328. https://doi.org/10.1016/S1053-8119(03)00085-5

Tanaka, J. W., & Simonyi, D. (2016). The “parts and wholes” of face recognition: a review of the literature. Quarterly Journal of Experimental Psychology, 218, 1–14. http://doi.org/10.1080/17470218.2016.1146780

Taubert, J., Apthorp, D., Aagten-Murphy, D., & Alais, D. (2011). The role of holistic processing in face perception: Evidence from the face inversion effect. Vision Research, 51(11), 1273–1278. http://doi.org/10.1016/j.visres.2011.04.002

Valentine, T. (1988). Upside-down faces: a review of the effect of inversion upon face recognition. British Journal of Psychology, 79(4), 471–491. http://doi.org/10.1111/j.2044-8295.1988.tb02747.x

Vida, M. D., Nestor, A., Plaut, D. C., & Behrmann, M. (2017). Spatiotemporal dynamics of similarity-based neural representations of facial identity. Proceedings of the National Academy of Sciences of the United States of America, 114(2), 388–393. http://doi.org/10.1073/pnas.1614763114

Winston, J. S., O’Doherty, J., & Dolan, R. J. (2003). Common and distinct neural responses during direct and incidental processing of multiple facial emotions. NeuroImage, 20(1), 84–97. https://doi.org/10.1016/S1053-8119(03)00303-3

Yin, R. K. (1969). Looking at upside-down faces. Journal of Experimental Psychology, 81(1), 141–145. http://doi.org/10.1037/h0027474

Zhang, H., Japee, S., Nolan, R., Chu, C., Liu, N., & Ungerleider, L. G. (2016). Face-selective regions differ in their ability to classify facial expressions. NeuroImage, 130, 77–90. http://doi.org/10.1016/j.neuroimage.2016.01.045

