## Supplementary material for "Spatio-temporal dynamics of face perception"

All supplementary analyses were conducted using same methods as the corresponding analyses in the paper, unless stated otherwise.

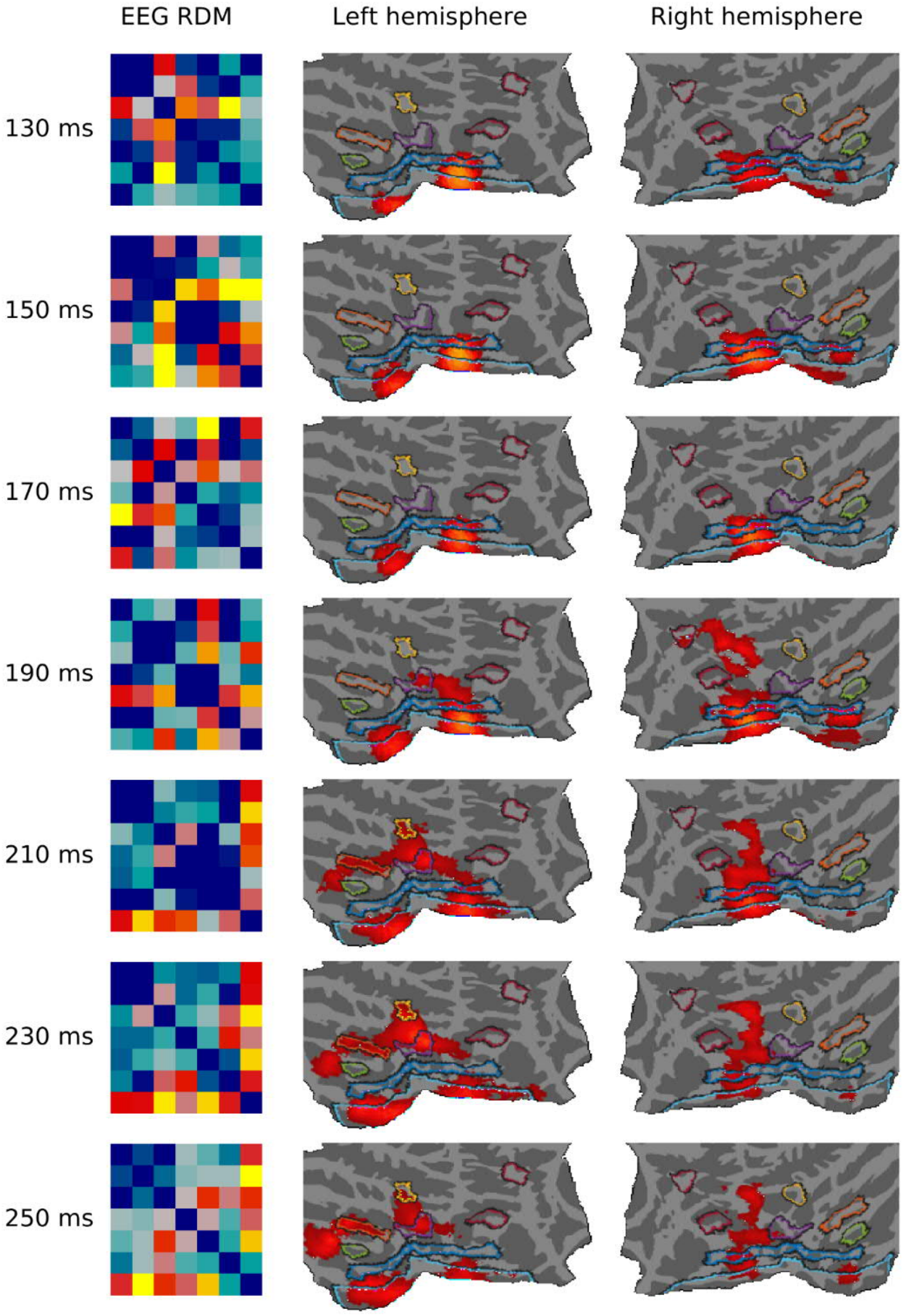


Supplementary Figure 1. Correlation between EEG and fMRI. EEG averaged (instead of fMRI). Voxel threshold pseudo-t=2.5. The correlations were mainly similar, although somewhat reduced, as in the original, fMRI-averaged analysis. With same voxel-level threshold (pseudo-t=3) as in the fMRI-averaged version used in the paper, similar correlations were found from V1, while the correlations in inferior temporal were found only at around 230 ms.

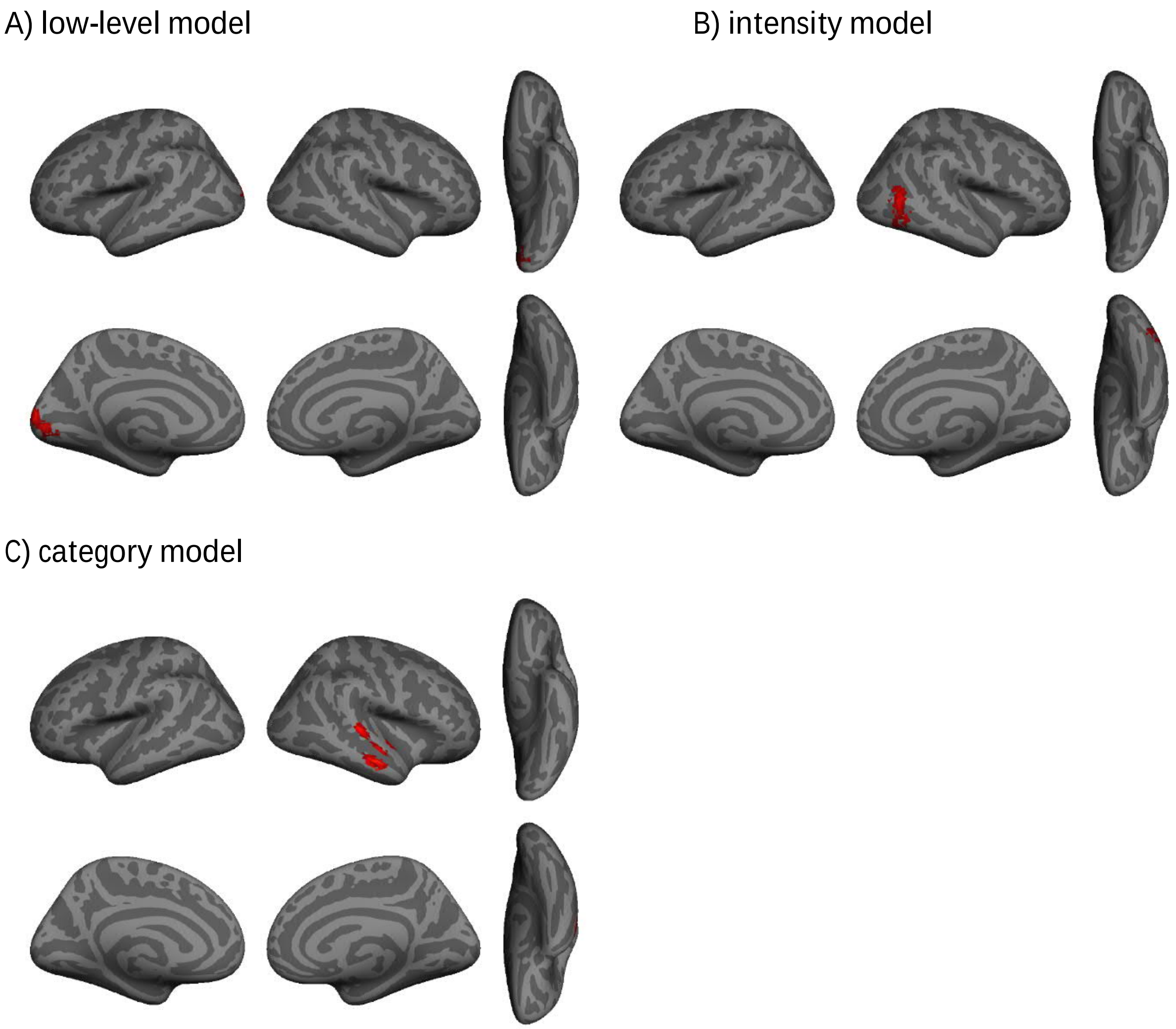


Supplementary Figure 2. fMRI model correlations without neutral faces. A) low-level model, B) intensity model, C) category model. Results for low-level and intensity models were similar as in the original analyses, while category model showed a significant cluster in anterior STS, not found when the neutral faces were included.

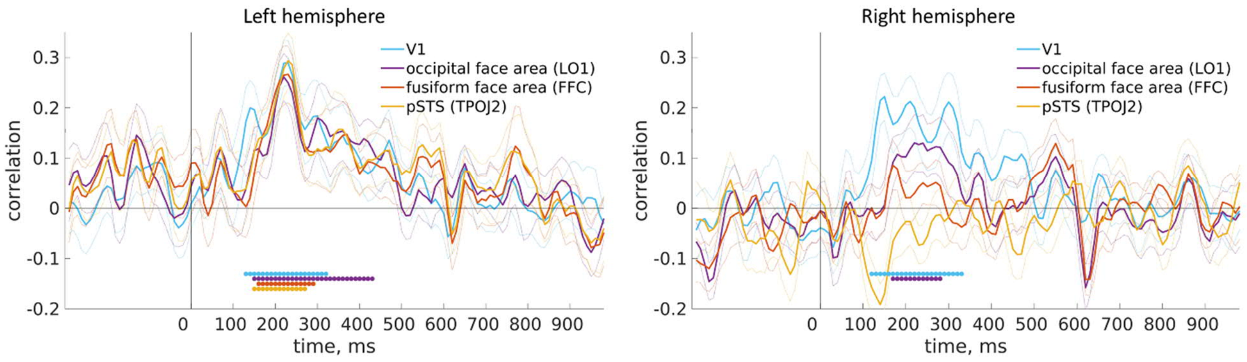


Supplementary Figure 3. ROI-EEG correlations with all voxels from each BALSA-ROI. Correlations were highly similar as when using five most informative voxels, except the right FFA did not correlate significantly with EEG.

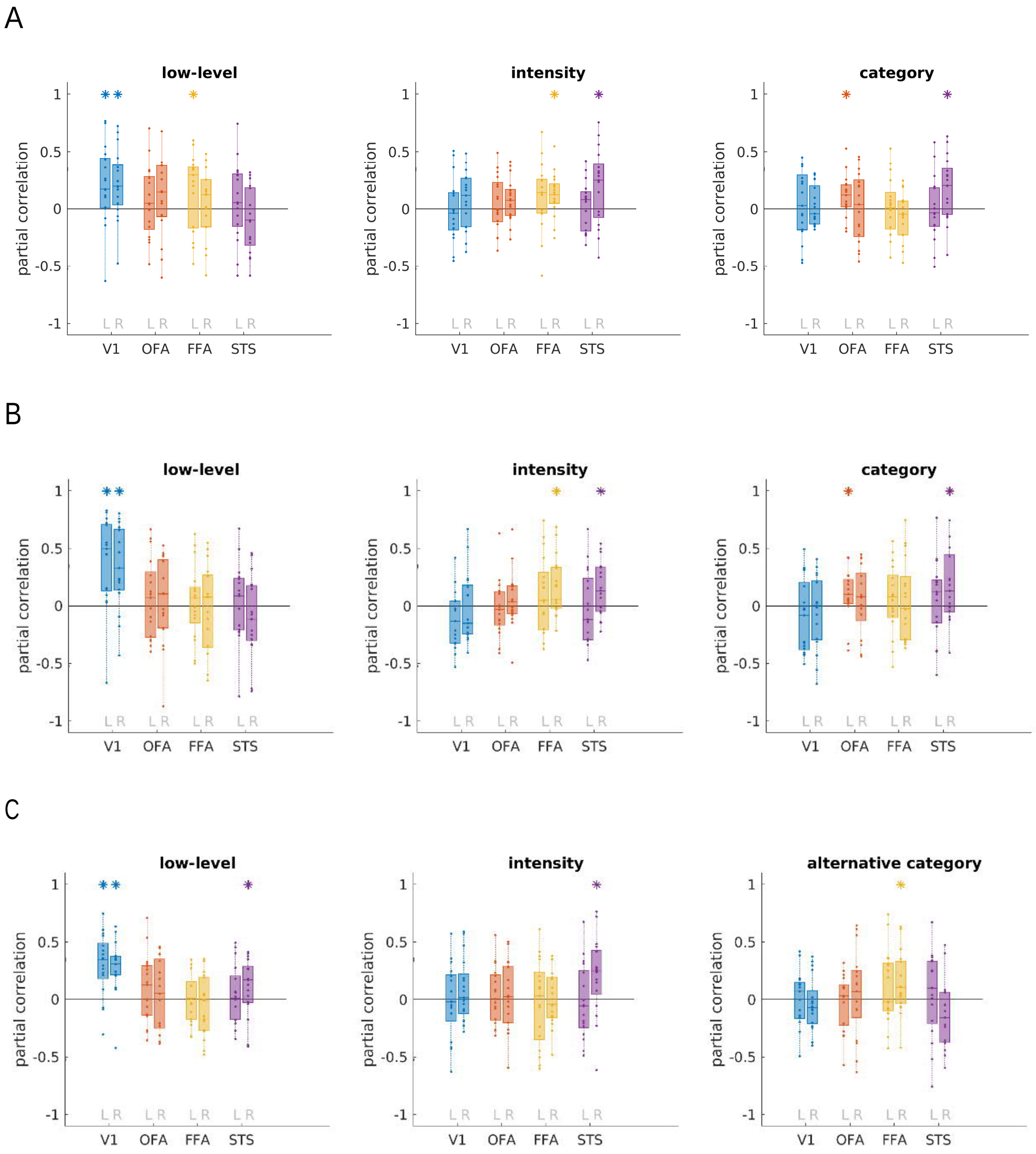


Supplementary Figure 4. fMRI model correlations in four different ROIs. A) Including all voxels (instead of five most informative) from each BALSA-ROI showed same significant ROIs as the original analysis. B) Removing neutral faces from the RDMs resulted in same significant ROIs except for left FFA in low-level model. C) Alternative category model, i.e., 50% morph equal to neutral. The main result of low-level model correlating with V1 and intensity model correlating with pSTS remained, while otherwise the results differed from the original category model. As the correlations are partial, the changing of one model affects the others as well. For wholebrain fMRI and EEG, the alternative category model did not show any significant correlations, as was found also with the original category model.
